# Proteolytic Cleavage by Matriptase Exacerbating Kidney Injury: a Novel Therapeutic Target

**DOI:** 10.1101/843607

**Authors:** Shota Ozawa, Masaya Matsubayashi, Hitoki Nanaura, Motoko Yanagita, Kiyoshi Mori, Katsuhiko Asanuma, Nobuyuki Kajiwara, Kazuyuki Hayashi, Hiroshi Ohashi, Masato Kasahara, Hideki Yokoi, Hiroaki Kataoka, Eiichiro Mori, Takahiko Nakagawa

**Affiliations:** TMK project at Medical Innovation Center, Kyoto University, Kyoto, Japan; Research Unit/Innovative Medical Science, Mitsubishi Tanabe Pharma Corporation, 3-2-10 Dosho-machi, Chuo-ku,Osaka, 541-8505, Japan; Department of Future Basic Medicine, Nara Medical University, 840 Shijo-Cho, Kashihara, Nara, 634-8521, Japan; Department of Nephrology, Graduate School of Medicine, Kyoto University, Shogoin-Kawahara-cho 54, Sakyo-ku, Kyoto, 606-8507, Japan; Institute for the Advanced Study of Human Biology (WPI-ASHBi), Kyoto University, Kyoto, 606-8501, Japan; Department of Molecular and Clinical Pharmacology, University of Shizuoka, 52-1 Yada,Suruga-ku,Shizuoka, 422-8526, Japan; Department of Nephrology, Chiba University, 1-8-1 Inohana, Chuo-ku, Chiba-shi, Chiba, 260-8677, JAPAN; Department of Nephrology, Ikeda City Hospital,3-1-18 Jonan, Ikeda, Osaka, 563-0025, Japan; Department of Pathology, Ikeda City Hospital,3-1-18 Jonan, Ikeda, Osaka, 563-0025, Japan; Institute for Clinical and Translational Science, Nara Medical University, 840 Shijo-Cho, Kashihara, Nara, 634-8521, Japan; Department of Pathology, University of Miyazaki, 5200 Kiyotakecho Kihara, Miyazaki-city, Miyazaki, 889-1692, Japan

**Keywords:** Chronic kidney disease, Matriptase, Hepatocyte growth factor activator inhibitor type 1, Podocin, Podocyte

## Abstract

Chronic kidney disease (CKD) is a progressive disease, and podocyte injury is a potential mechanism. We found that Matriptase was activated at podocytes in CKD patients and mice while a Matriptase inhibitor slowed the progression of mouse kidney disease. The mechanism could be accounted for by an imbalance favoring Matriptase over its cognate inhibitor, hepatocyte growth factor activator inhibitor type 1 (HAI-1), as conditional depletion of HAI-1 in podocytes accelerates podocyte injury. Intriguingly, the N-terminal of Podocin (Podocin-N), as a consequence of Matriptase cleavage of Podocin, translocates to nucleoli. These results suggest that aberrant activation of Matriptase would cause podocyte injury, and a targeting Matriptase could be a novel therapeutic strategy for CKD patients.

**Significant statement:** Chronic kidney disease (CKD) is a progressive disease. If podocytes, which are specialized cells of the kidney glomerulus that wrap around capillaries, are injured, kidney injury is exacerbated. Thus, a therapeutic strategy to addressing CKD would be to block podocyte injury. The present study provides evidence that Matriptase cleaves Podocin, a component of podocyte slit membrane, and the N-terminal of podocin translocates to nucleoli, causing kidney injury. Our findings show that the N-terminal of Podocin plays an efficient role for cell fate in podocytes. In addition, the inhibition of Matriptase would be a potential therapeutic target for CKD.

## Introduction

Chronic kidney disease (CKD) has come to be recognized as a global public health problem (1). An increasing number of CKD patients end up suffering from end stage renal disease, requiring renal hemodialysis, and loading huge financial burden on the shoulder of their home countries. Developing specific treatments for curing CKD are warranted.

The glomerular filtration unit consists of three layers: fenestrated endothelial cells, the glomerular basement membrane, and podocytes (2). Podocytes, a major cell type of renal glomerulus, are terminally differentiated, and interdigitate with adjacent podocytes to form glomerular filtration barrier (3). Interdigitated foot processes of podocytes form a 40-nm-wide cell junction that is composed of several proteins called the slit diaphragm (SD).

Accumulating evidence demonstrate that intracellular proteolytic processing is required to maintain physiological function in podocytes while aberrant activation causes cellular injury (4). Podocyte injury is often initiated by SD disruption, but its precise mechanism remains unclear. Recently, it was shown that a non-specific serine protease inhibitor, camostat mesilate (CM), slowed the progression of renal disease due to the protection of podocytes (5), suggesting that serine proteases should be involved. However, the specific type of enzyme involved in this process also remains unknown.

## Results

Transcriptome analysis using glomeruli of Adriamycin (ADR) nephropathy mice identified 15 serine proteases, which were highly upregulated (>2.0 fold) in glomeruli of mice with ADR nephropathy. This is in comparison with control mice at 6 weeks after administration (Fig.1A, Supplemental Table 1). Among these factors, Matriptase (also known as suppressor of tumorigenicity 14 protein, ST14), only a type II transmembrane serine protease (TTSP), is expressed at the epithelial cell junctions in a wide variety of tissues (6). We then examined the expressions of other TTPS and found that Matriptase was most highly induced at glomeruli in ADR mice (Fig.1 B-C). Subsequent analyses demonstrated that Matriptase was induced at mRNA and protein levels in podocytes of ADR mice at day 42 (Fig. 2A-C, Fig.1B-C). Glomerular Matriptase protein was also significantly higher in patients with clinical proteinuria due to diabetic nephropathy or membranous nephropathy, but not in subjects with IgA nephropathy who had no clinical proteinuria (Fig. 2D).

**Fig. 1.**
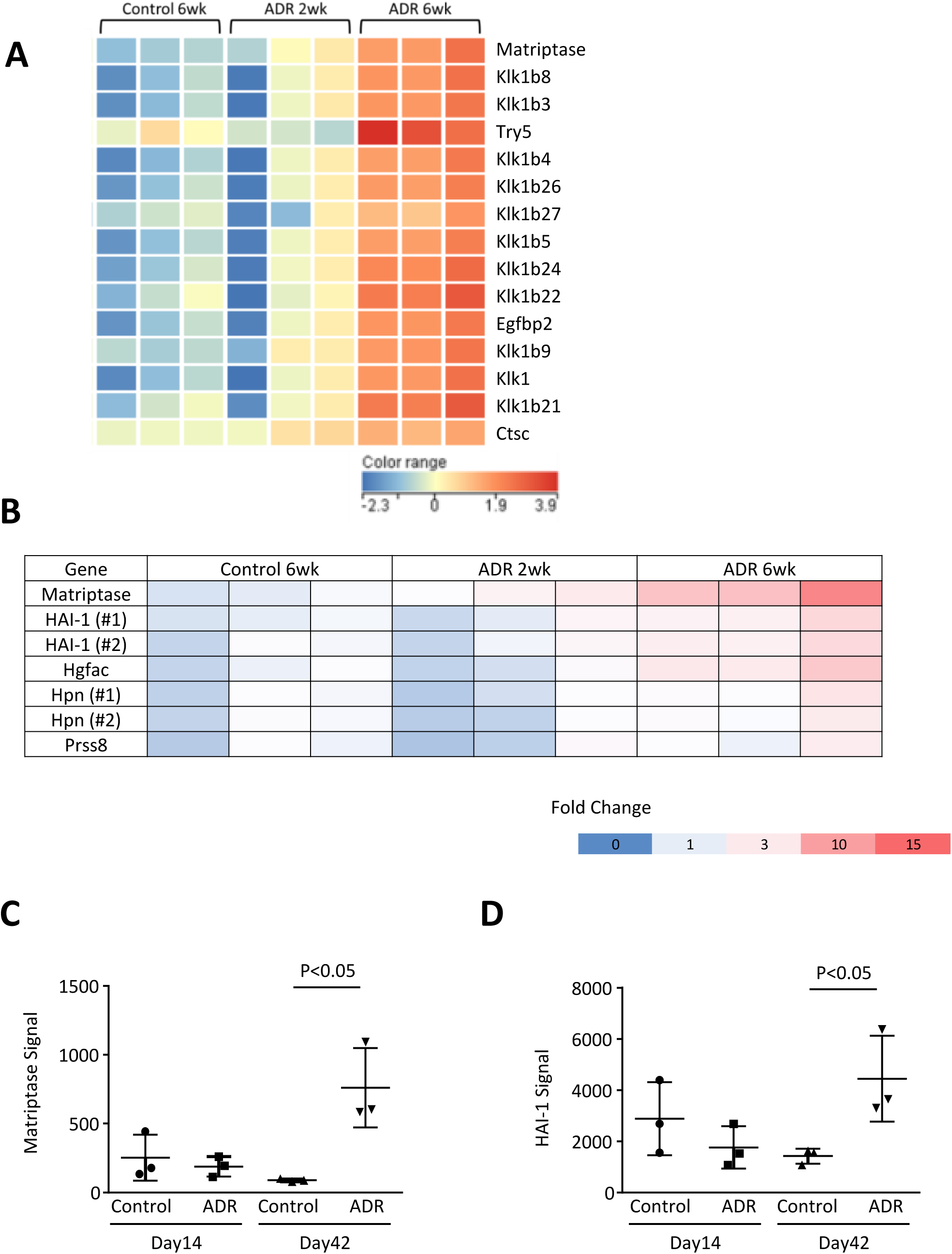
Transcriptome analysis. (A) Microarray was performed with glomerular RNA of control mice (N=3), and ADR nephropathy mice at 2 weeks (N=3) and at 6 weeks (N=3). Statistical significance (p<0.05) was calculated using moderated student’s t-test followed by Benjamini-Hochberg false discovery rate correction on GeneSpring GX. Fold change cut-off of > 2 lead to extract 357 probes in ADR nephropathy mice at 6 weeks compared with control mice. Among those genes, 15 genes are categorized as serine protease while only Matriptase is a type II membrane anchored serine protease. Heatmap shows that all serine proteases are upregulated at 42 days in glomerulus of mice with ADR nephropathy. (B) Among 7 probes, which are categorized as membrane anchored serine proteinase and its cognate inhibitors, only Matriptase and HAI-1 are significantly upregulated in ADR-nephropathy mice compared with control mice (B, C, D). Hgfac, hepatocyte growth factor activator; Hpn, hepsin; Prss8, serine protease 8

**Fig. 2.**
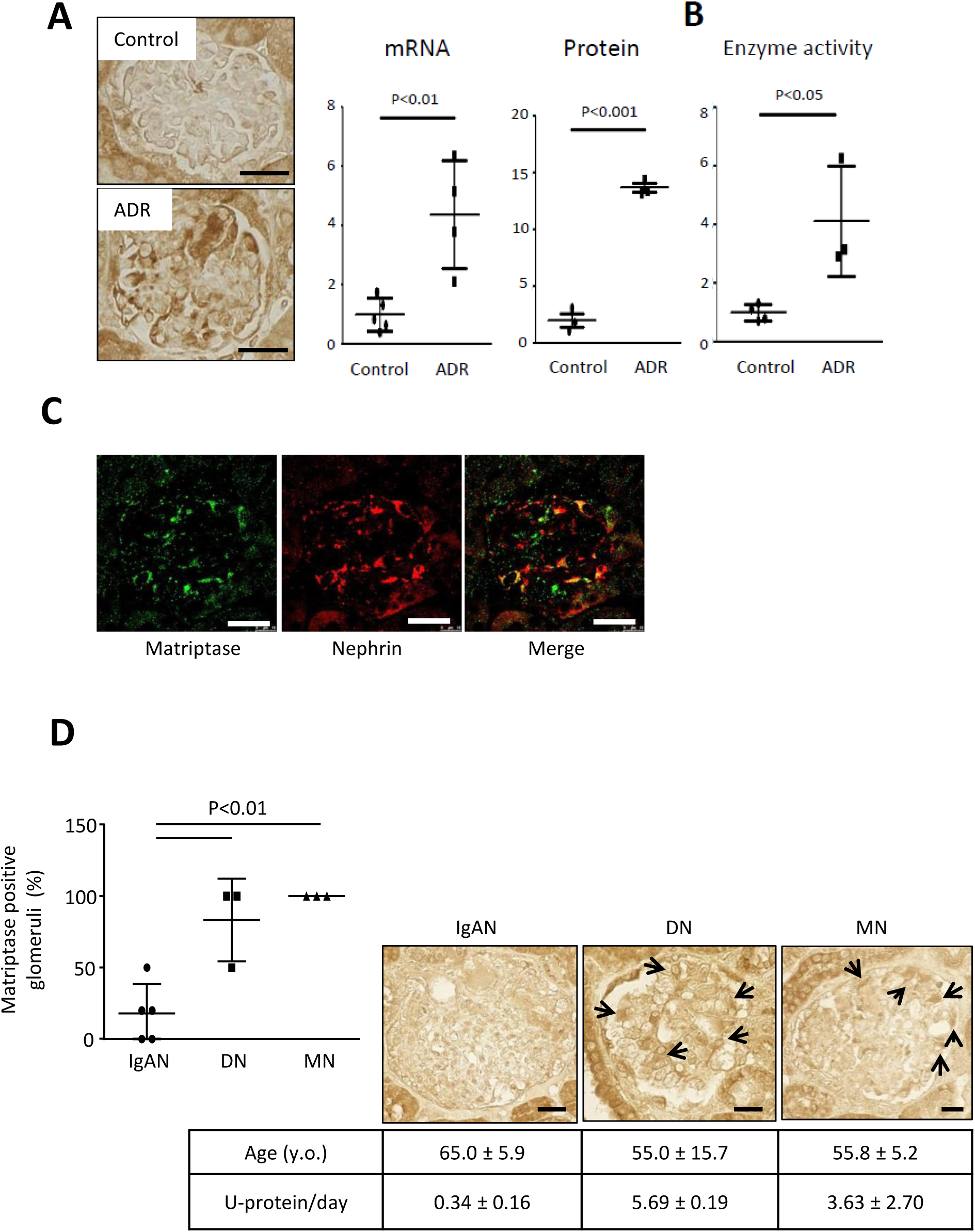
Matriptase is induced in podocytes of mice with ADR nephropathy and patients with clinical proteinuria. (A) Immunohistochemistry shows an increase in glomerular Matriptase protein expression, and mRNA expression and protein of Matriptase are significantly higher in glomeruli of ADR-induced nephropathy mice. (B) Glomerular serine protease activity is enhanced in ADR mice compared with control mice. (C) Matriptase (green) is co-localized (yellow) with Nephrin (red) in glomerulus of mouse with ADR nephropathy. (D) Matriptase protein expression is observed in human glomerulus of diabetic nephropathy (DN) and membranous nephropathy (MN) while it is negative in that of IgA nephropathy (IgAN). Bar: 20 μm

Matriptase is tightly regulated by its cognate inhibitor HAI-1 (also known as SPINT1), a Kunitz-type serine protease inhibitor (7). HAI-1 mRNA expression was significantly induced in the glomeruli of ADR mice compared to that of wild type mice (Fig. 1B, D). Subsequently another type of mouse model with podocyte injury (8, 9), streptozotocin-induced diabetic endothelial nitric oxide synthase knockout (diabetic eNOS-KO) mice, was examined for subsequent transcriptome analysis using isolated glomeruli (Supplemental Table 2). Among 13 candidate genes (Fig. S2A), HAI-1 level was significantly upregulated in the podocytes (confirmed by immunohistochemistry) and in serum of diabetic eNOS-KO mice (Fig. 3A-C, Fig. S2B). A transcriptome analysis showed that glomerular Matriptase mRNA expression was highly upregulated by 5.3 times in diabetic eNOS-KO mice compared to eNOS-KO mice (Fig.S2C-D). Analyses of these two independent mouse models suggest that the Matriptase/HAI-1 pathway plays a key role in podocyte injury and therefore, it could be a potential therapeutic target for kidney disease.

**Fig. 3.**
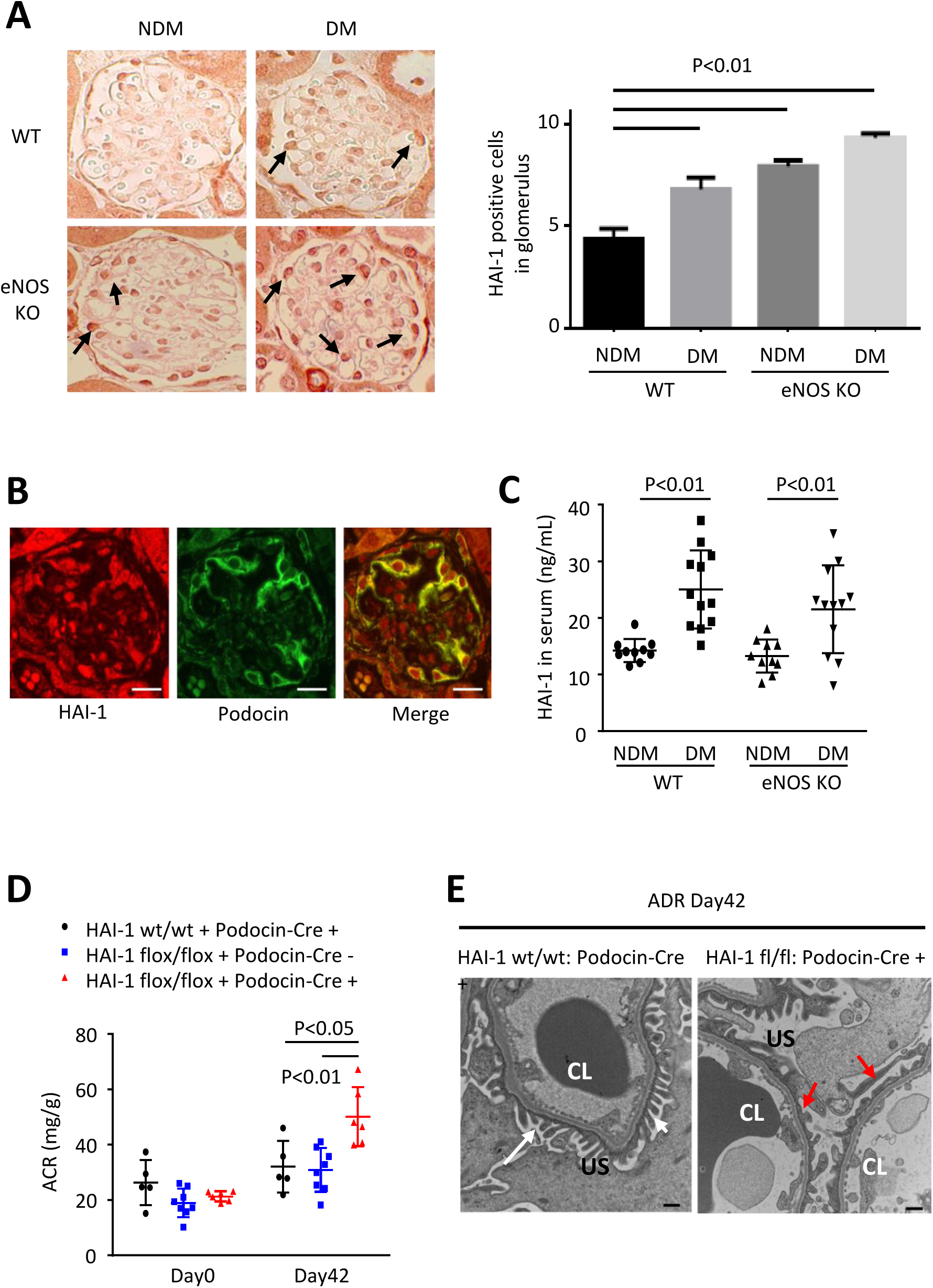
Role of HAI-1 in diabetic eNOS-KO mice and mice with ADR nephropathy. (A) The number of HAI-1 positive cells (arrows) increases in glomeruli of STZ-induced diabetic mice and eNOS-KO mice, and it is further higher in diabetic eNOS-KO mice than in any other groups. (B) HAI-1 partially co-localized with Podocin. (Bar: 20μm). (C) HAI-1 level in serum is also higher in diabetic (DM) wild type and diabetic eNOS-KO mice compared with non-diabetic (NDM) wild type and NDM eNOS-KO mice. (D) 42 days after 10.5 mg/kg ADR was injected into HAI-1 wt/wt: Podocin-Cre (+) mice (N=5), HAI-1 flox/flox: Podocin-Cre (-) mice (N=8), and HAI-1 flox/flox: Podocin-Cre (+) mice (N=6), urinary albumin/creatinine ratio (ACR) was examined. (E) Compared to foot process (white arrows) in HAI-1 wt/wt: Podocin-Cre (+) mice, the effacement (red arrows) of podocytes in HAI-1 flox/flox: Podocin-Cre (+) injected with ADR is observed under Transmission Electron Microscopy. (Bar: 500nm). CL, capillary lumen; US, urinary space.

Matriptase and HAI-1 are co-expressed in many epithelial cells (10), and the regulation of Matriptase by HAI-1 is required for epidermal integrity (11). In turn, the imbalance favoring Matriptase over HAI-1 contributes to various diseases (12-15). According to the Nephroseq database (https://www.nephroseq.org/), HAI-1 is induced (1.5 fold), along with an increase in Matriptase expression (2.0 fold), in the podocytes of diabetic nephropathy. This indicates that enhancing Matriptase activation over HAI-1 could be a potential mechanism. Consistent with these observations, the podocyte-specific depletion of HAI-1 deteriorated podocyte injury in ADR nephropathy (Fig. 3D and 3E, Fig S3A and S3B). The podocyte injury could be accounted for by aberrant Matriptase activation due to the absence of HAI-1 in the podocyte.

To develop a therapeutic approach to nephropathy, we subsequently examined whether the selective inhibition of Matriptase could protect kidneys from disease progression. We examined two types of serine protease inhibitors toward Matriptase activity; a selective peptide-mimetic inhibitor of Matriptase (IN-1) (16) and a synthetic non-specific serine protease inhibitor Nafamostat mesilate (NM), which ameliorates rat kidney disease (5). It was found that the two compounds inhibited Matriptase activity in a dose response manner, and inhibitory effects were nearly equipotent, with half-maximal inhibitory concentration of IN-1 (IC_50 IN-1_) = 1.3 nM versus IC_50 NM_ = 0.86 nM (Fig. 4A). However, inhibition assays toward thrombin activity showed that IC_50_ was 2.8 μM for IN-1, whereas it was 99 nM for NM (Fig. 4B), suggesting that IN-1 is more specific to Matriptase than NM. To examine the therapeutic effect of IN-1, this compound was applied to BALB/c mice with ADR nephropathy upon 15 days after ADR treatment. We observed that Matriptase inhibitor IN-1 slowed the progression of ADR nephropathy with blocking podocyte injury and ameliorated albuminuria (Fig. 4C-F). These results indicate that blocking Matriptase could be a potential therapeutic approach against chronic kidney diseases.

**Fig. 4.**
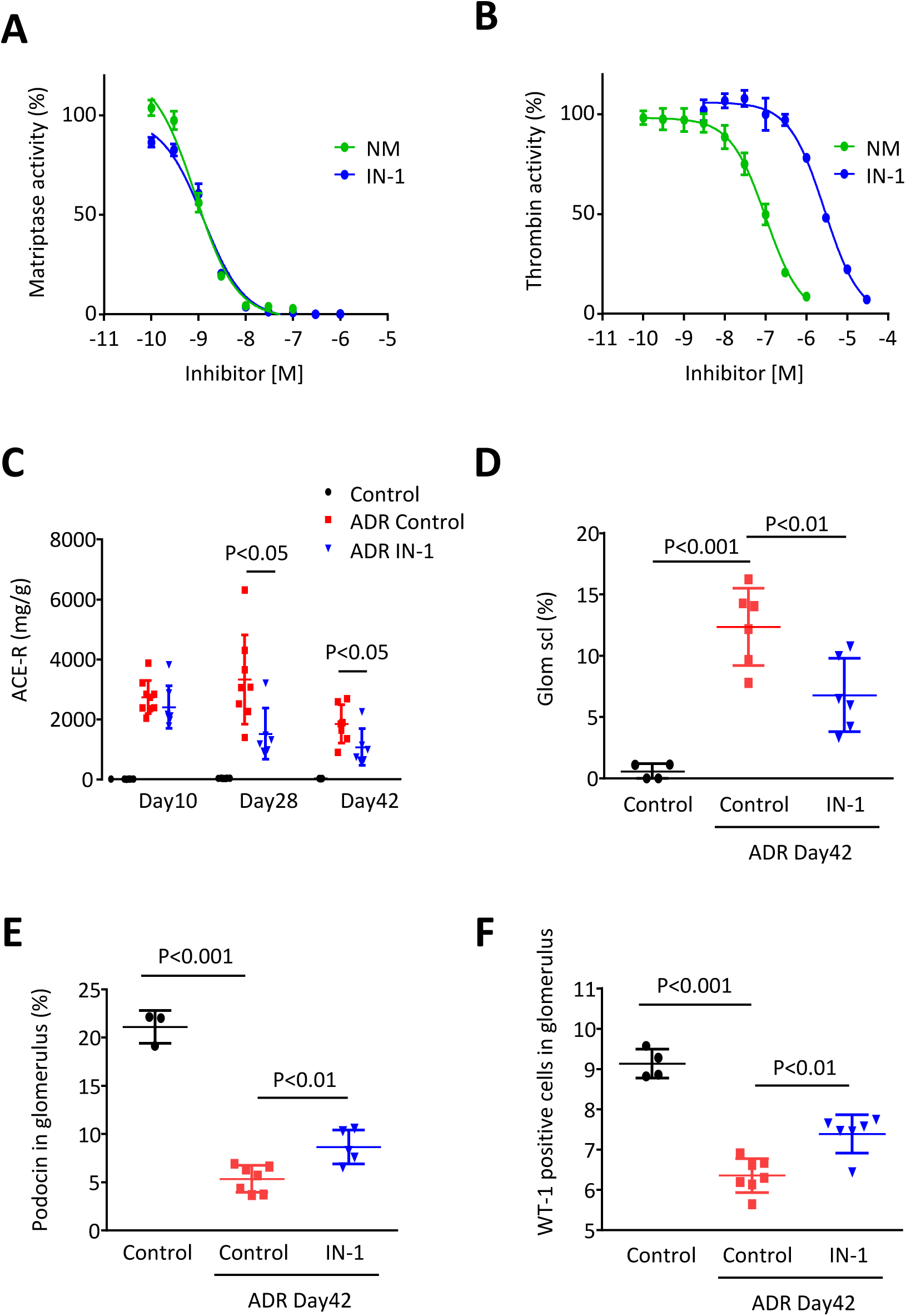
Pharmacological inhibition of Matriptase slows the progression of ADR nephropathy in mice. (A) IC_50_ of IN-1 for Matriptase activity is 1.3 nM in a protease activity assay whereas that of Nafamostat mesilate (NM) is 0.86 nM. (B) IC_50_ of IN-1 for Thrombin activity is 2.8 μM whereas that of NM is 99 nM. (C) Urinary albumin/creatinine ratio (ACR) is higher in mice with ADR nephropathy, but it is significantly suppressed by the injection of 10 mg/kg IN-1 three times a week on day 28 and on day 42. (D) Glomerular sclerosis is higher in mice with ADR nephropathy compared to normal control mice. However, it is significantly reduced by IN-1 treatment on day 42. (E, F) Podocin positive area and the number of WT positive cells in glomerulus are significantly lower by ADR treatment, but such detrimental effect of ADR is slightly but significantly ameliorated by the chronic treatment with IN-1 on day 42.

Next, we sought to identify the substrates of Matriptase in podocytes. Interestingly, Matriptase was found to directly cleave mouse Podocin (mPodocin) (Fig. S5A, S5I), but not with other slit membrane proteins; Claudin5, β-Catenin or Nephrin (Fig. S5B-D). Enzymatic dead G827R Matriptase mutant failed to cleave mPodocin (Fig. 5A), and mPodocin was cleaved at the membrane but not in the cytoplasm (Fig. 5B). A mutagenesis on mPodocin identified R50 as the specific cleavage site of Matriptase, and not at R36, R45, or R54 (Fig. 5C, Fig. S5H-I). Matriptase cleavage of mPodocin was blocked by co-expression of wild type as well as extracellular domain of HAI-1 (Fig. 5D). We also identified the fragmented cleaved mPodocin in the urine of mice with ADR nephropathy (Fig. S5E), suggesting that Matriptase cleaves Podocin in *in vivo*. Together, these results suggest that mPodocin cleaved by Matriptase might be a potential mechanism underlying the progression of podocyte injury.

**Fig. 5.**
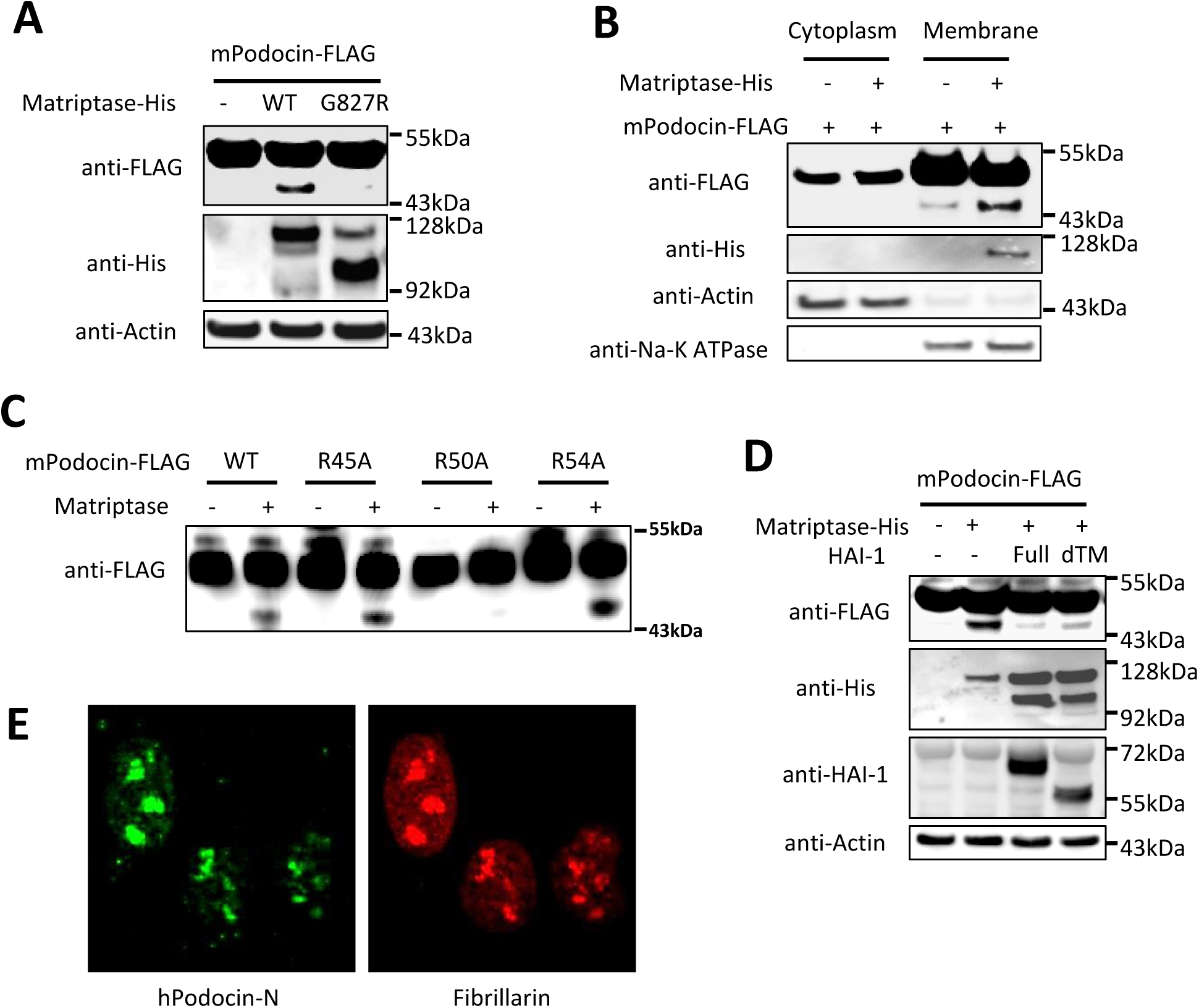
Matriptase cleaves Podocin at R50, which is blocked by HAI-1. (A) While wild type (WT) of Matriptase cleaves Podocin and produces its small fragment, the inactive form of Matriptase (G827R) fails to do it in HEK293 cells overexpressing mPodocin-FLAG and either WT-Matriptase or G827R Matriptase mutant. (B) Matriptase-induced Podocin cleavage occurs at the membrane fraction, but not in the cytoplasm of HEK293 cells overexpressing Matriptase-His and mPodocin-FLAG. (C) Matriptase cleavage of Podocin R50A mutant, as opposed to wild type, R45A and R54A mutants, is declined using in vitro cleavage assay. (D) Full length of HAI-1 or secreted type of HAI-1 (deleted transmembrane region (dTM)) blocks the cleavage of Podocin in response to Matriptase in HEK293 cells. (E) N-terminal fragment of human Podocin tagged with 3xFLAG is translocated into nucleus, particularly in nucleoli of U2OS cells.

Exogenous expression of mPodocin N-terminal fragment (mPodocin-N) (2-50aa) showed the mPodocin-N as being localized in the nucleus, while full length mPodocin and ΔN-mPodocin (51-385aa) remained at the plasma membrane (Fig. S6A and S6B). Human Podocin (hPodocin) was also cleaved by Matriptase (Fig. S5G), and N-terminal fragment of hPodocin (hPodocin-N) migrated to the nucleus and bound nucleoli (Fig 5E).

## Discussion

Podocytes have been found to produce other types of proteases, matrix metalloproteases MMP-2 and -9, all of which are believed to have significant roles within the glomerulus as they degrade type IV collagen, a major protein of the glomerular basement membrane (17). It has also been observed that altered MMPs/tissue inhibitors of MMPs (TIMPs) balance may lead to increased extracellular matrix deposition or excessive degradation activity – a finding with critical implications for glomerular diseases (2). The upregulation of MMP-9 expression can be found in several glomerular diseases accompanied by proteinuria in human patients (18, 19) and diabetic nephropathy murine model (20). Podocytes express CD40 and CD154, the ligand of CD40, to induce MMP-9 expression in an autocrine fashion (21). In various diseases, coagulation proteases also contribute to tissue injuries, including cancer progression and cardiovascular diseases. These injuries are mediated by protease-activated receptors (PARs) (22, 23), a family of G protein-coupled receptors consisting of four members (PAR1-PAR4). Consistent with findings from an earlier study which suggested that PARs play a role in kidney disease progression (24), PAR-1 has indeed been found to contribute to the development of podocyte and glomerular injuries (25).

It is important to realize that individual proteases induce divergent signaling pathways that lead to differential functional consequences (26). These studies imply that the strategy to targeting all the proteases via non-specific protease inhibitor may not always be beneficial; the targeting of specific protease(s) by specific inhibitor(s) may be a better approach. Although proteases play key roles in the maintenance of renal filtration barrier and there are indications to suggest that serine protease inhibition is involved in kidney disease progression (5), the precise mechanism(s) underlying the progression of kidney disease, in which serine proteases are involved, as well as the target serine protease(s), remain unknown and to be identified.

In this study, we showed that HAI-1 was essential to the protection of podocytes from nephropathy by blocking of Matriptase-mediated cleavage of Podocin. We also demonstrated that a transmembrane serine protease Matriptase acted as a potential therapeutic target, findings which imply a new class of selective peptide-mimetic inhibitor of Matriptase can be applied as a potential drug for the treatment of chronic kidney diseases. Although careful pre-clinical studies are required before applying the Matriptase inhibitor in clinical practice, our finding that NM, which is commonly used for the treatment of pancreatitis and disseminated intravascular coagulation (27), also protects the kidney from disease progression, might give some reassurance that the inhibition of Matriptase is a safe and promising approach.

In conclusion, we identified Matriptase as a potential cause for podocyte injury. The aberrantly activated enzyme beyond the suppression by HAI-1 potently cleaves Podocin and disturbs podocyte integrity, leading to renal injury. Most surprisingly, Podocin-N migrates to the nucleus and binds nucleoli, indicating its roles in physiological and active cell death process.

## Materials and Methods

### Animal Study

All animal experiments were performed in accordance with either the Animal Experimentation Committee of Kyoto University or Tanabe R&D Service Co., Ltd. (Osaka, Japan), or both. Male C57BL/6J-Nos3tm1nc (eNOS-KO) mice at 7 weeks of age were purchased from the Jackson Laboratory (Bar Harbor, ME). Mice with diabetic nephropathy were developed as described previously (8). Diabetes was induced with intraperitoneal injection of 50 mg/kg/day streptozotocin (STZ) for five consecutive days. Diabetes was defined as non-fasting blood glucose > 250 mg/dl using a blood glucose meter. Mice were fed a standard laboratory chow ad libitum. Mean blood pressure was measured using CODA Multi-Channel, Computerized, Non-Invasive Blood Pressure System (Kent Scientific Corporation, Torrington, CT) while blood glucose level was determined by GLUCOCARD MyDIA (Arkray, Edina, MN). Urine was collected overnight using metabolic cages. At 14 weeks of age, all mice were sacrificed.

For ADR nephropathy model, ADR at a dose of 10.5 mg/kg body weight was injected via the tail vein of male BALB/c mice at 8 weeks of age. Matriptase inhibitor (IN-1) at a dose of 10 mg/kg body weight was intraperitoneally injected to mice with ADR nephropathy three times a week for 4 weeks. Urine was collected overnight using metabolic cages. At 14 weeks of age, all mice were sacrificed. Matriptase inhibitor (IN-1) was synthesized (WuXi AppTec Co., Ltd., Shanghai, China) as previously described (16). HAI-1 flox/flox mice were kindly provided from Dr. Hiroaki Kataoka (28). Establishment of Podocin-CreERT2 [Podocin-Cre (+)] mice was reported previously (29). HAI-1 flox/flox: Podocin-Cre (+) mice were produced by crossing HAI-1 flox/flox mice and Podocin-Cre mice. The primers for genotyping Podocin-Cre were TTTGCCTGCATTACCGGTCGATGCAAC and TGCCCCTGTTTCACTATCCAGGTTACGGA. The primers used for genotyping HAI-1 flox were ACCACTGGCTCATTTGGTGTTGGC and TGAAGCCTGGCCACTTCCTGATG.

To induce functional Cre protein, 150 mg/kg Tamoxifen for 3 consecutive days was injected intraperitoneally at the age of 8 weeks. ADR at a dose of 10.5 mg/kg body weight was injected via the tail vein of male mice at 10 weeks of age. At 16 weeks of age, all mice were sacrificed.

### Molecular Analysis

Urinary albumin concentration was measured using Albuwell M (Exocell, Philadelphia, PA) or MICROFLUORAL Microalbumin Test (PROGEN, Heidelberg, Germany). Urine creatinine was measured using LabAssay creatinine (Wako, Osaka, Japan). Serum HAI-1 concentration was measured using mouse HAI-1 ELISA kit (R&D Systems, Minneapolis, MN).

### Isolation of Glomeruli

Mice were perfused with 8 × 10^7^ Dynabeads (Life technology, Waltham, MA) diluted in 40 mL of Hank’s Balanced Salt Solution through the heart under anesthesia. Kidneys were isolated and digested in collagenase solution (1 mg/ml collagenase A and 100 U/ml deoxyribonuclease I) for 30min at 37°C, and then glomeruli containing Dynabeads were gathered by a magnetic particle concentrator. During the procedure, kidney tissues were kept at 4°C except for the collagenase digestion. Finally, glomerular RNA was purified with RNeasy Mini Kit (Qiagen, Chatsworth, CA). For measuring protease activity in glomeruli, glomeruli were lysed with RIPA buffer (Santa Cruz Biotechnology Inc., Dallas, TX).

### Microarray Analysis

Glomerular RNA was extracted and purified with RNeasy Mini Kit (Qiagen, Chatsworth, CA). RNA quality was assessed with Agilent 2100 Bioanalyzer (Agilent Technologies, Santa Clara, CA). In accordance with the Agilent Technologies protocol, all samples were processed and hybridized to SurePrint G3 Mouse Gene Expression 8 × 60 K (Agilent Technologies). Fluorescence was detected using Agilent DNA Microarray Scanner. The data was analyzed with GeneSpring GX (Agilent Technologies).

### Histological Analysis

Formalin-fixed, paraffin-embedded sections (2 μm) were stained with periodic acid-Schiff reagent (PAS) for the light microscopy. Glomeruli (50-100 per kidney) were examined on coronal sections to evaluate the degrees of glomerulosclerosis. Glomerulosclerosis was defined as obstruction of the capillary lumen caused by mesangial expansion or collapsed capillaries.

### Immunohistochemistry and Immunofluorescence

Following deparaffinization, the formalin-fixed, paraffin-embedded section samples were incubated with 3% H_2_O_2_ for 20min to inactivate endogenous peroxidase activity. Samples were incubated with citrate buffer (pH 6.0) for retrievals of antigens for Podocin, Nephrin and WT-1. The sections were subsequently incubated with primary antibodies. The following antibodies were used: anti Nephrin antibody (R&D Systems), anti-HAI-1 antibody (R&D Systems), anti-WT-1 antibody (Santa Cruz Biotechnology Inc), and anti-Podocin antibody (Sigma Aldrich, St. Louis, MO). For Matriptase antibody, anti-serum was collected from rabbits immunized with peptide corresponding mouse and human Matriptase sequence (single letter code, CAQRNKPGVYTRLP). After reaction with primary antibody, sections were incubated with ImmPRESS Reagent Kit (Vector Labs, Burlingame, CA) for immunohistochemistry or incubated with Alexa Fluor conjugated secondary antibodies (Invitrogen, Carlsbad, CA) for immunofluorescence. Positive areas were measured using MetaMorph (Molecular Devices, Sunnyvale, CA). After permeabilization for 10 min, fixed U2OS was incubated in blocking solution [5% normal goat serum+0.1% Triton X-100 in PBS] and then incubated with primary antibodies diluted in blocking solution overnight at 4 °C. Cells were incubated with Alexa Fluor conjugated secondary antibodies diluted in the blocking solution for 1h at room temperature and mounted on slide glass with VECTASHIELD Mounting Medium with DAPI (Vector Laboratories). The following antibodies were used: anti-FLAG M2 monoclonal Antibody (Sigma Aldrich), anti-Fibrillarin (Cell Signaling Technology).

Samples were analyzed using Olympus confocal microscope Fluoview FV-1000 under 100x objective lens.

### Quantitative Polymerase Chain Reaction (qPCR)

cDNA was synthesized from purified RNA using ReverTra Ace qPCR RT Kit (TOYOBO, Osaka, Japan). Real-time PCR was performed using a Step One Plus thermal cycler (Applied Biosystems, Waltham, MA). Amount of PCR products was normalized with 18S rRNA mRNA. Following primers were used:

18S rRNA

AGGGGAGAGCGGGTAAGAGA (forward (FW)) and

GGACAGGACTAGGCGGAACA (reverse (RV));

Matriptase TCCCTCAGAGCCAGAAGTGT (FW) and

ACAGTCCGTCTTCCCATCAC (RV);

Podocin

GTGTCCAAAGCCATCCAGTT (FW) and

GTCTTTGTGCCTCAGCTTCC (RV);

Synaptopodin

TTCCGAGTGGCATCCTTAAGTC (FW) and

GCTGCTGCTTGGTAGGTTCA (RV).

### Cell Culture

HEK293 was used for transfection of plasmids with FuGENE HD Transfection reagent (Promega, Tokyo, Japan). RIPA buffer was used for obtaining proteins from whole cells. For separating cytoplasm and membrane proteins, subcellular protein fractionation kit (Thermo Fisher Scientific) was used. U2OS was seeded in cover glass on 6-well plate (Falcon) with DMEM complete medium. Next day, transfection was performed using FuGENE HD Transfection reagent. After 24h, cells were fixed with 4% paraformaldehyde.

### Plasmid Constructs

A full length cDNA of mouse Podocin was cloned into pFLAG-CMV-6a (Sigma Aldrich, St.Louis,MO). cDNA of mutant Podocin (R45A, R50A and R54A) was generated by PCR and cloned into pFLAG-CMV-6a. A full length cDNA of Matriptase constructed in pcDNA3.1-V5-His (Thermo Fisher Scientific) was provided by Dr. Hiroaki Kataoka. Matriptase mutant (G827R) was generated by PCR and cloned into pcDNA3.1-V5-His. A full length of cDNA of HAI-1 and the deletion type of transmembrane region cloned into pCIneo were also provided by Dr. Kataoka. A human Podocin-N fragment cDNA with 3xFLAG was amplified by PCR using a human kidney cDNA in Human MTC Panel I (TaKaRa) as a template. A 3xFLAG Podocin-N was cloned into pcDNA3.1(+).

### In vitro cleavage assay

Cell lysate from HEK293 transfected with FLAG-Podocin (WT, R45A, R50 and R54A) was immobilized on anti-FLAG beads (Sigma-Aldrich). The beads were washed with IP buffer (1%–2% Triton-X, 150 mM NaCl, 50 mM Tris-HCl [pH 7.5]). Samples eluted with FLAG peptide was used as substrate for Matriptase. 15 ng/mL Recombinant Mouse Matriptase (R&D Systems) and the substrate were incubated in assay buffer including 50 mM Tris-HCL (pH 8.5), 50 mM NaCl and 0.01% Tween20 for 2hr.

### Western Blotting

Cell lysate was separated by sodium dodecyl sulfate polyacrylamide gel electrophoresis and electrotransferred onto polyvinylidene fluoride membranes. After overnight incubation with primary antibody at 4°C, the membranes were incubated with HPR-conjugated anti-mouse (GE Healthcare, Buckinghamshire, England) or anti-rabbit (GE Healthcare) secondary antibody for 1hr at room temperature, followed by addition of ECL prime (GE Healthcare) to detect bands using Image Quant LAS4000mini (GE Healthcare). Primary antibodies were used as follows; anti FLAG antibody (Sigma Aldrich), anti His antibody (MBL, Nagoya, Japan), anti Actin antibody (Sigma Aldrich), anti NA-K ATPase antibody (Cell Signaling, Danvers, MA) and anti-HAI-1 antibody (R&D Systems). All blots were run under the same experimental conditions. Each image was obtained at single time point and was not combined into a single image.

### Protease Activity Assay

Matriptase or Thrombin activity was assessed by measuring 7-Amino-4-methylcoumarin (AMC) release from synthetic substrates. For Matriptase activity, 25 μM substrate (Boc-QAC-AMC) (Enzo Life Sciences, Farmingdale, NY),100 pg/μL Recombinant Mouse Matriptase (R&D Systems) in assay buffer including 50 mM Tris-HCl (pH 8.5), 50 mM NaCl and 0.01% Tween20. For Thrombin activity, 60 mM Unit Thrombin (SIGMA-Aldrich) was used instead of Matriptase. Nafamostat mesilate (Sigma Aldrich) or Matriptase inhibitor (IN-1) was added 10min before adding substrate. For both assays, the released fluorescence (Ex: 380 nm, Em: 460 nM) was measured using an EnVision (Perkin Elmer) at a certain time.

### Transmission Electron Microscopy

Kidneys were fixed with 4% PFA and 2% glutaraldehyde with 0.1 M phosphate buffer (PB) at 4°C overnight. After post-fixation with 1% Osmium Tetroxide in 0.1 M PB for 2 hr, samples were dehydrated in ethanol and propylene oxide. Then, the samples were penetrated in propylene oxide and Epon after polymerization in pure Epon. Ultrathin sections were cut with an ultramicrotome. The sections were stained in uranyl acetate and lead citrate. The grids were examined with a transmission electron microscope (H-7650; Hitachi).

### Analysis of human kidney specimens

All of the human specimens were procured and analyzed after informed consent and with approval of the Ethics Committee of Ikeda City Hospital and Kyoto University. Tissue samples were obtained from diagnostic renal biopsies performed at Ikeda City Hospital. We investigated samples from patients who had been diagnosed with IgA nephropathy (N=5), diabetic nephropathy (N=3) and membranous nephropathy (N=3).

### Nephroseq database analysis

A kidney transcriptomics data repository Nephroseq (www.nephroseq.org, University of Michigan) was used to analyze deposited various data sets. The expression of HAI-1, HAI-2 and Matriptase was checked in the diabetic nephropathy dataset of comparison of Healthy Living with diabetic nephropathy (30). We checked the gene expressions of glomeruli (threshold; p<0.05, fold change>1.5) and tubulointerstitium (threshold; none).

### Statistical Analysis

All values are presented as mean ± SD. Statistical analysis was perfomred with ANOVA using Tukey’s method to compare groups or two-tailed t test. A level of p<0.05 was considered statistically significant.

## Acknowledgements

The authors thank Prof. Pierre Chambon for the CreERT2 plasmid, and Keren-Happuch E Fan Fen for critical reading of the manuscript. This work was supported by grants from TMK Project, JSPS KAKENHI [26460274 to T.N., JP17H07031 to E.M., JP18H06202 to H.N.], SUZUKEN CO., LTD. to T.N., MSD Lifescience Foundation to T.N. and E.M., Novartis Research Grants to T.N., E.M., FUJI YAKUHIN CO., LTD. to T.N., E.M., Takeda Science Foundation to E.M., Kanzawa Medical Research Foundation to E.M., Uehara Memorial Foundation to E.M., Nakatomi Foundation to E.M., Konica Minolta Science and Technology Foundation to E.M., Naito Foundation to E.M., Mochida Memorial Foundation for Medical and Pharmaceutical Research to E.M., SENSHIN Medical Research Foundation to E.M., Terumo Foundation for Life Sciences and Arts to E.M., Nara Kidney Disease Research Foundation to E.M., and by unrestricted funds provided to E.M. from Dr. Taichi Noda (KTX Corp., Aichi, Japan) and Dr. Yasuhiro Horii (Koseikai, Nara, Japan).

