## Supplemental Figure for "Proteolytic Cleavage by Matriptase Exacerbating Kidney Injury: a Novel Therapeutic Target"

**Supplemental Figure legends**

**Fig S1.** In order to further examine the specific mechanism, we first investigated serine protease activities in the kidney using the Adriamycin (ADR) murine nephropathy model. ADR at a dose of 10.5 mg/kg body weight was injected via the tail vein of male BALB/c mice at 8 weeks of age. At 10 or 14 weeks of age, all mice were sacrificed. Glomerular RNA was collected and evaluated as described in Materials and Methods. (A) ADR-induced nephropathy mice show a significant increase in urinary albumin excretion on day 42. Podocyte injury was also confirmed by significant reductions in mRNA expressions of Podocin, Synaptopodin, and Wilms tumor protein (WT-1: a transcription factor for the specific activation of a glomerular differentiation program in renal precursors (31)) in mice with ADR-induced nephropathy compared with those in control mice (B-D). ADR was also induced glomerulosclerosis in mice, in which Podocin and Nephrin were down-regulated and pathologically dislocated in the glomeruli (E).

**Fig S2.** (A) Samples were glomerular RNA extracted from mice shown in Table S2. After microarray analysis, several steps were applied to extract key genes. Final 13 genes are shown. (B) Urinary albumin/creatinine ratio in wild type and eNOS KO mice with/without diabetes in Figure 3A-C is shown. (C) (D) Fold changes between two groups and gene expression data about HAI-1 and TTSPs included in Figure 1B are shown.
NDM, non-diabetes; DM, diabetes

**Fig S3.** The mouse with podocyte specific HAI-1 knockout mice. (A) In immunohistochemistry, HAI-1 flox/flox: Podocin-Cre (+) mice shows a reduction in HAI-1 expression after injection of Tamoxifen. Red arrows show HAI-1 expression in podocytes. (B) HAI-1 positive area per glomerulus significantly decreases in HAI-1 flox/flox: Podocin-Cre (+) after tamoxifen treatment. Bar: 20μm

**Fig S4.** (A) We treated ADR-induced nephropathy mice with NM from day 15 to day 42. (B) PAS staining shows severe glomerular sclerosis in mice with ADR nephropathy. Treatment with Nafamostat mesilate (NM) ameliorated glomerular injury in ADR mice. (C) Immunohistochemistry shows that Podocin expression is reduced in ADR mice, but it is ameliorated by NM treatment. (D) ADR mice develop albuminuria, which is suppressed by nafamostat mesilate. Data of ACR from Control and ADR Control is the same as those used in Figure 4C.

**Fig S5.** (A) Co-expression of FLAG-Podocin and Matriptase in HEK293 gives rise to the cleaved form of Podocin. (B, C) A full length cDNA of Claudin 5 or β-Catenin was cloned into pFLAG-CMV-6a. Claudin5 or β-Catenin was overexpressed with Matriptase in HEK293 cells. No effect of Matriptase on Claudin 5 and β-Catenin are shown. (D) HeLa cells were infected by adenovirus for expressing Nephrin-DsRed and transfected with Matriptase. No effect of Matriptase on Nephrin protein is shown. (E) Urinary protein concentrated with ITSIPrep Total Protein Isolation Kit (ITSI BIOSCIENCE, Johnstown, PA) contains the cleaved Podocin in ADR nephropathy mice. Glomerular (Glom) protein and anti-albumin antibody (Exocell) were used. (F) Matriptase cleaves Podocin in dose-dependent manner in HEK293. (G) Human podocin-tagged with myc extracted from HEK293 cells is cleaved by Matriptase in dose-dependent manner. (H) N-terminal 64 amino acids of mouse Podocin are shown highlighting Arginine, which could be a cleavage site for Matriptase, in red. (I) Podocin (Full length) and deleted type of Podocin (△1-36) are cleaved by Matriptase, producing △N-mPodocin.

**Fig S6.** (A) Transfection of FLAG-mPodocin or FLAG-mPodocin(2-50) or FLAG-Podocin(51-385) in MDCK are shown. Anti-FLAG antibody detects each protein as a result of western blotting. (B) Red signal indicates location of FLAG-tagged podocin. Podocin-N (2-50aa) is localized in the nucleus whereas full length Podocin and ΔN-Podocin (51-385) are stayed in cytosol.

**Table S1.** General characteristics of mice with ADR nephropathy.

**Table S2.** General characteristics of mice with diabetic eNOSKO mice.

### Fig S1

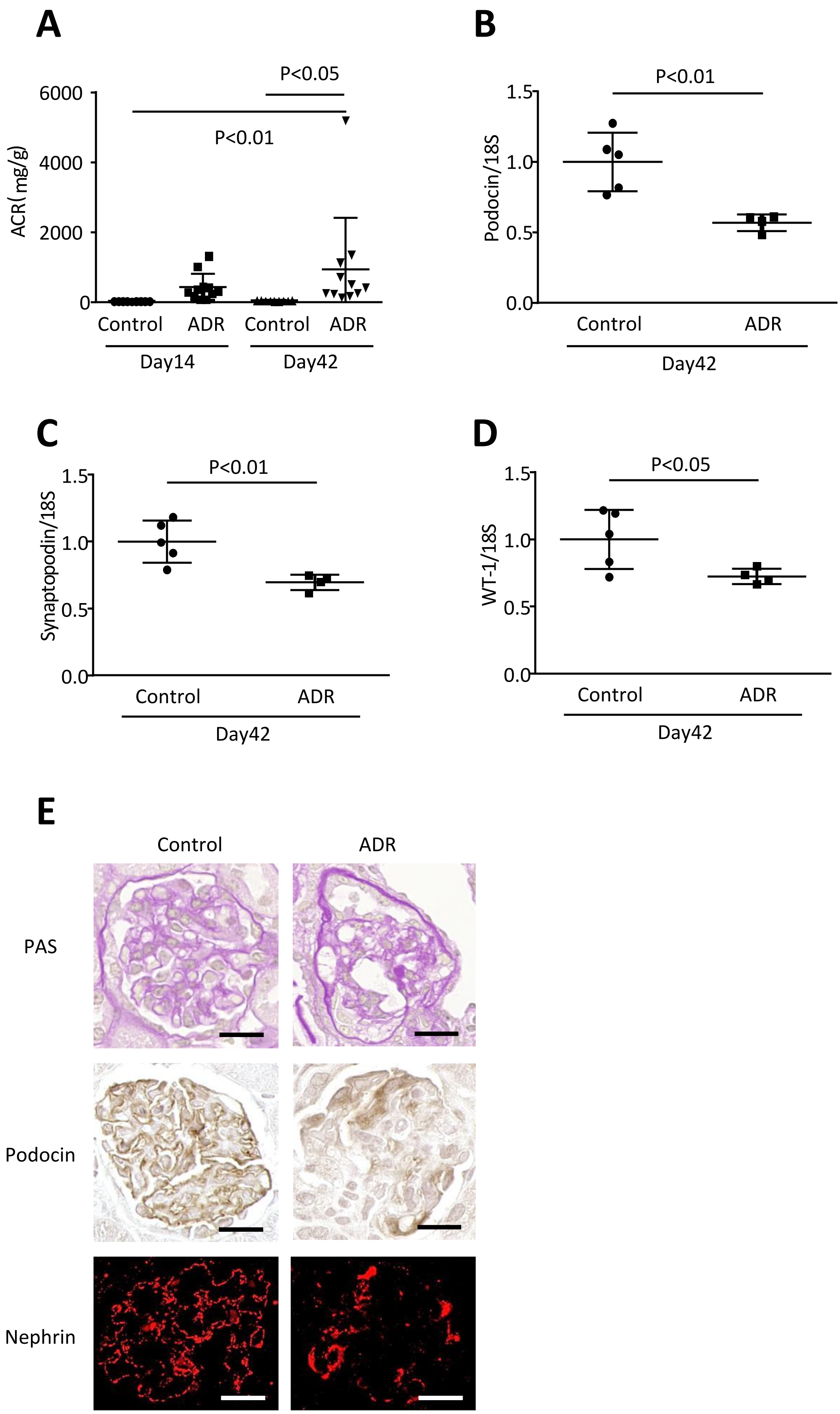

A
P<0.05
P<0.01
ACR( mg/g)
6000
4000
2000
0
Control
ADR
Control
ADR
Day14
Day42
B
P<0.01
Podocin/18S
1.5
1.0
0.5
0.0
Control
ADR
Day42
C
P<0.01
Synaptopodin/18S
1.5
1.0
0.5
0.0
Control
ADR
Day42
D
P<0.05
WT-1/18S
1.5
1.0
0.5
0.0
Control
ADR
Day42
E
Control
ADR
PAS
Podocin
Nephrin

### Fig S2

## A

Group1: NDM-eNOSKO 14 weeks old
Group2: DM-eNOSKO 14 weeks old (diabetic for 6 weeks)

55861 probes

↓ Group2/Group1>2-fold

3086 probes

↓ >100 in raw data of Group2

938 probes

↓ Group2/Group1>6.5-fold

110 probes

↓ Desirable pattern in Nephromine

13 genes;
Spint1, Tspan1, Lad1, Wfdc2, Tmprss2, Pigr, Krt23, Fabp3, Tox3, Sfrp1, Gsta2, Pgam2, Spp1

## B

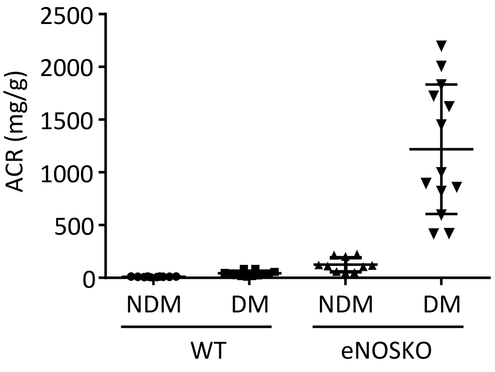


## C

Microarray

| | | Fold change Group2 / Group1 |
|---|---|---|
| | Spint1 | 6.5 |
| Serine protease activity | St14 (Matriptase) | 5.3 |
| | Prss8 (Prostasin) | 3.7 (13.9) |
| | Hpn (Hepsin) | 4.3 |
| | Hgfac | 12.1 |

## D

| GeneSymbol | normalized G1 | normalized G2 | Flags G1 | Flags G2 | raw G1 | raw G2 |
|---|---|---|---|---|---|---|
| Spint1 | -3.173 | -0.570 | Not Detected | Detected | 32.136 | 219.756 |
| Spint1 | -2.000 | 0.705 | Detected | Detected | 72.483 | 531.887 |

| | | | | | | |
|---|---|---|---|---|---|---|
| St14 (Matriptase) | -1.966 | 0.444 | Detected | Detected | 74.204 | 443.724 |
| Prss8 | -1.381 | 0.535 | Detected | Detected | 111.262 | 472.583 |
| Prss8 | -3.425 | 0.443 | Not Detected | Detected | 26.989 | 443.332 |
| Hpn | 1.320 | 3.386 | Detected | Detected | 723.701 | 3408.850 |
| Hgfac | -4.980 | -1.401 | Not Detected | Detected | 9.181 | 123.557 |

### Fig S3

**A**

HAI-1 wt/wt
Podocin-Cre +

HAI-1 flox/flox
Podocin-Cre +

HAI-1

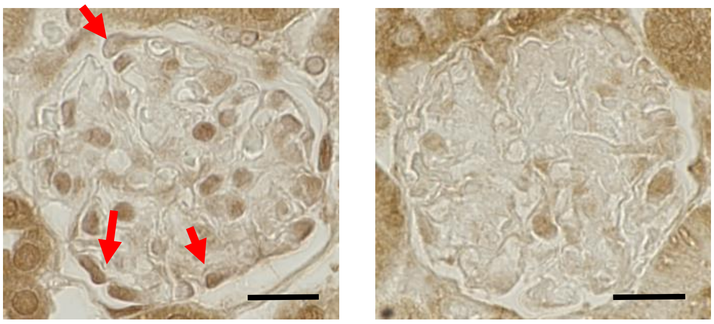

**B**

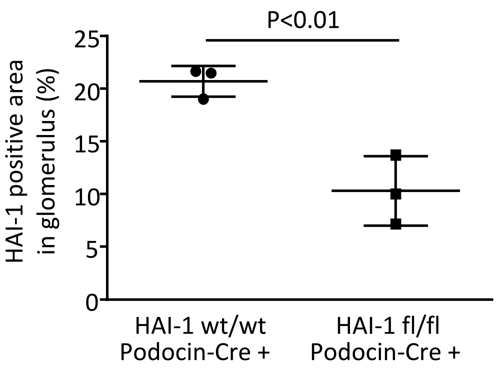

**Fig S4**

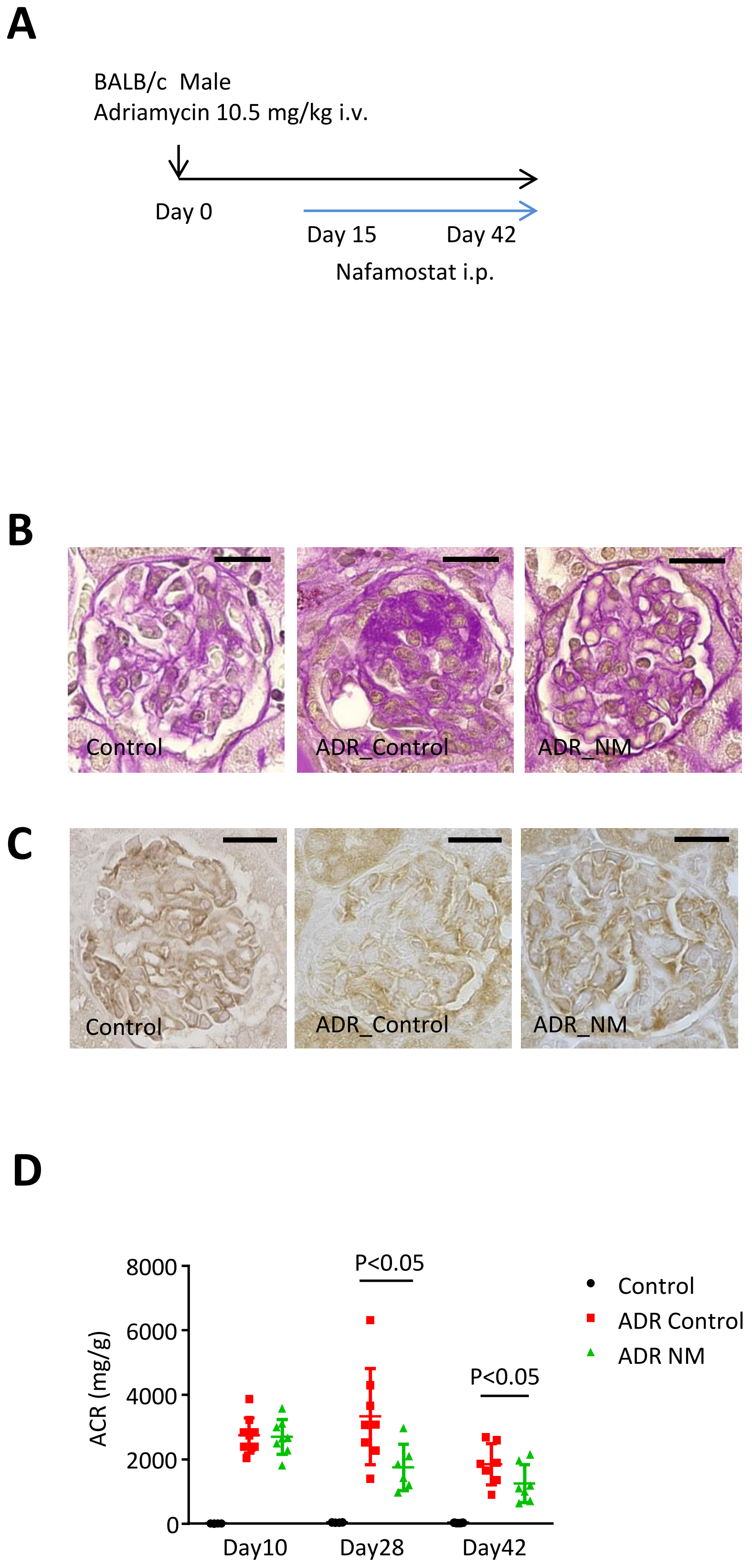
A
BALB/c Male
Adriamycin 10.5 mg/kg i.v.
Day 0
Day 15
Day 42
Nafamostat i.p.
B
Control
ADR_Control
ADR_NM
C
Control
ADR_Control
ADR_NM
D
ACR (mg/g)
8000
6000
4000
2000
0
Day10
Day28
Day42
P<0.05
P<0.05
Control
ADR Control
ADR NM

#### Fig S5

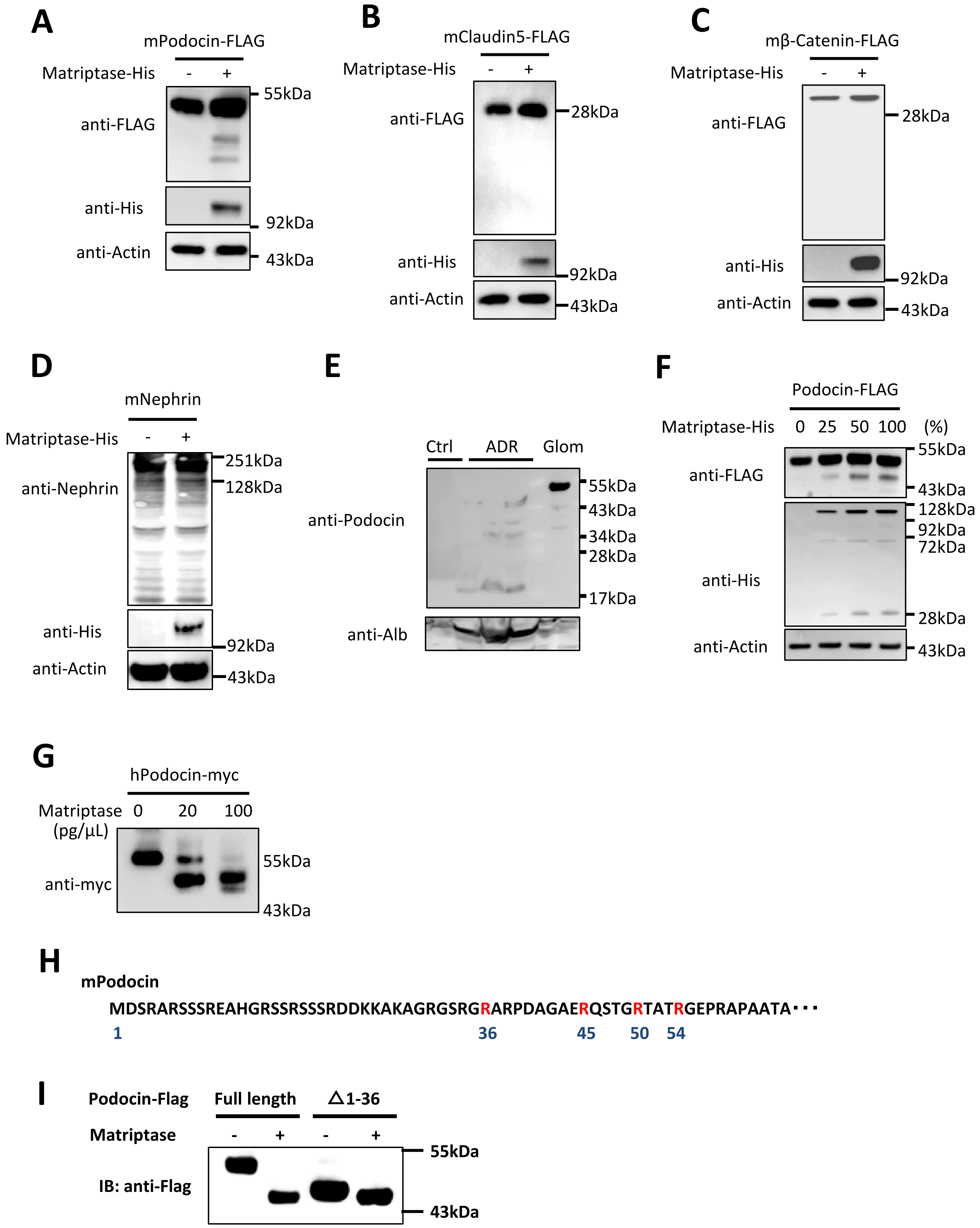

A
mPodocin-FLAG
Matriptase-His - +
anti-FLAG
anti-His
anti-Actin
55kDa
92kDa
43kDa
B
mClaudin5-FLAG
Matriptase-His - +
anti-FLAG
anti-His
anti-Actin
28kDa
92kDa
43kDa
C
mβ-Catenin-FLAG
Matriptase-His - +
anti-FLAG
anti-His
anti-Actin
28kDa
92kDa
43kDa
D
mNephrin
Matriptase-His - +
anti-Nephrin
anti-His
anti-Actin
251kDa
128kDa
92kDa
43kDa
E
Ctrl ADR Glom
anti-Podocin
anti-Alb
55kDa
43kDa
34kDa
28kDa
17kDa
F
Podocin-FLAG
Matriptase-His 0 25 50 100 (%)
anti-FLAG
anti-His
anti-Actin
55kDa
43kDa
128kDa
92kDa
72kDa
28kDa
43kDa
G
hPodocin-myc
Matriptase (pg/μL) 0 20 100
anti-myc
55kDa
43kDa
H
mPodocin
MDSRARSSSREAHGRSSRSSSRDDKKAKAGRGSRGRARPDAGAERQSTGRTATRGEPRAPAATA···
1 36 45 50 54
I
Podocin-Flag Full length △1-36
Matriptase - + - +
IB: anti-Flag
55kDa
43kDa

### Fig S6

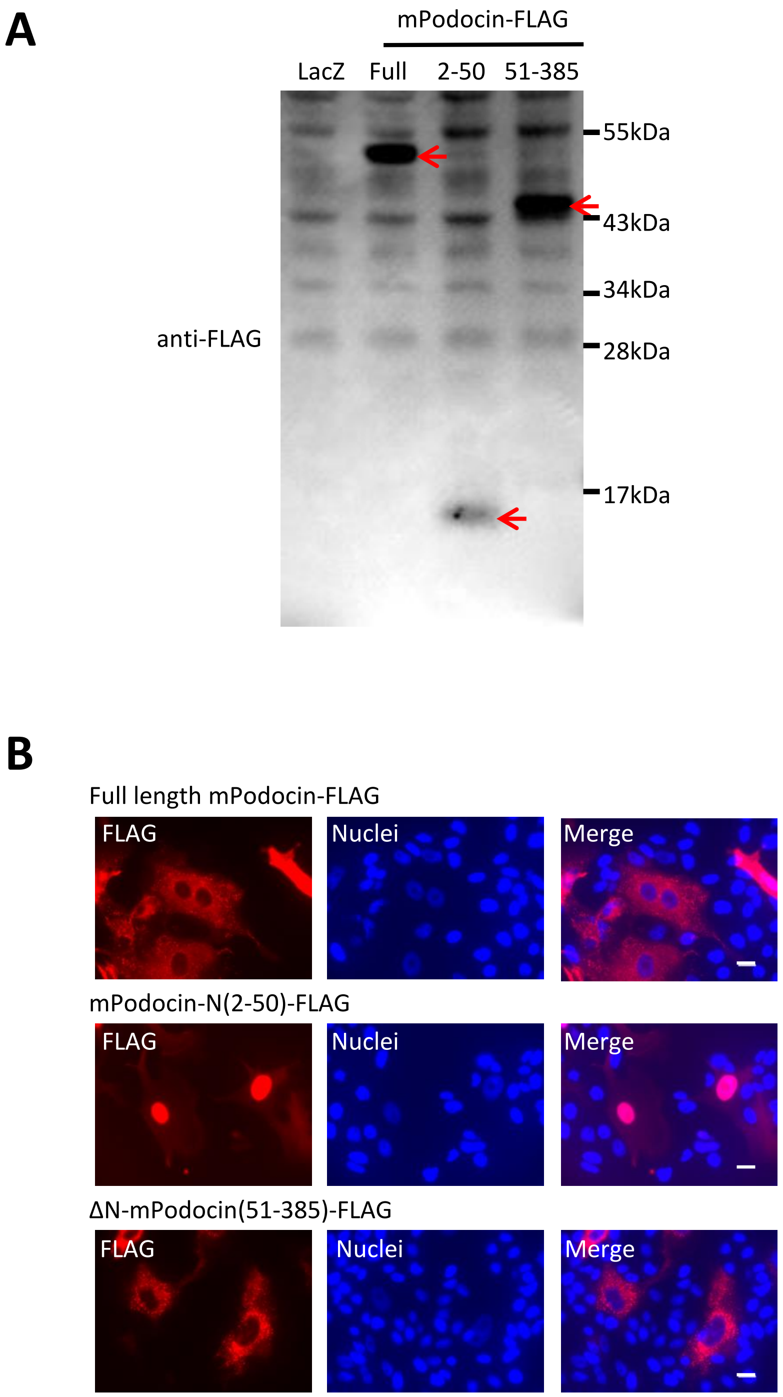
A
mPodocin-FLAG
LacZ
Full
2-50
51-385
55kDa
43kDa
34kDa
28kDa
17kDa
anti-FLAG
B
Full length mPodocin-FLAG
FLAG
Nuclei
Merge
mPodocin-N(2-50)-FLAG
FLAG
Nuclei
Merge
ΔN-mPodocin(51-385)-FLAG
FLAG
Nuclei
Merge

### Table S1

| | Control | | | ADR | | |
|---|---|---|---|---|---|---|
| Age | 8 week | 10 week | 14 week | 8 week | 10 week | 14 week |
| Body weight (g) | 23.9±1.2 | 24.6±1.3 | 27.7±1.6 | 23.0±0.7 | 23.8±1.0 | 24.6±1.3 |
| Albumin creatine ratio (mg/g) | ND | 16.7±5.6 | 31.4±7.8 | ND | 599.4±246.5 | 1067.7±325.0 |

ADR, adriamycin; ND, not done

### Table S2

| | eNOSKO mice | | | | | |
|---|---|---|---|---|---|---|
| | NDM | | | DM | | |
| Age | 8 week | 10 week | 14 week | 8 week | 10 week | 14 week |
| Body weight (g) | 21.0±1.0 | 26.1±0.7 | 26.3±1.2 | 19.3±1.0 | 22.7±4.0 | 24.0±3.2 |
| Blood glucose (mg/dl) | ND | 121.3±19.1 | 105.3±1.5 | ND | 402.0±179.9 | 506.7±10.0 |
| Systolic blood pressure (mmHg) | ND | 116.5±12.3 | 128.2±8.8 | ND | 126.9±18.7 | 125.5±13.6 |
| Diastolic blood pressure (mmHg) | ND | 92.1±10.9 | 105.0±10.1 | ND | 103.2±15.1 | 99.5±10.8 |
| Urine albumin (μg/day) | 60.5±5.3 | 96.5±30.8 | 73.3±43.0 | 83.8±45.2 | 474.6±592.3 | 223.4±104.6 |
| Albumin creatine ratio (mg/g) | 263.2±55.5 | 415.2±69.6 | 231.5±162.4 | 314.3±163.4 | 1505.5±1792.0 | 573.2±287.9 |

NDM, non-diabetic mellitus; DM, diabetic mellitus; eNOSKO, eNOS knock-out; ND, not done
